# How pirate phage interferes with helper phage: Comparison of the two distinct strategies

**DOI:** 10.1101/576587

**Authors:** Namiko Mitarai

## Abstract

Pirate phages use the structural proteins encoded by unrelated helper phages to propagate. The best-studied example is the pirate P4 and helper P2 of coliphages, and it has been known that the *Staphylococcus aureus* pathogenicity islands (SaPIs) that can encode virulence factors act as pirate phages, too. When alone in the host, the pirate phages act as a prophage, but when the helper phage gene is also in the same host cell, the pirate phage has ability to exploit the helper phages structural proteins to produce pirate phage particles and spread, interfering with the helper phage production. The known helper phages in these systems are temperate phages. Interestingly, the interference of the pirate phage to the helper phage occurs in a different manner between the SaPI-helper system and the P4-P2 system. SaPIs cannot lyse a helper lysogen upon infection, while when a helper phage lyse a SaPI lysogen, most of the phage particles produced are the SaPI particles. On the contrary, in the P4-P2 system, a pirate phage P4 can lyse a helper P2 lysogen to produce mostly the P4 particles, while when P2 phage lyses a P4 lysogen, most of the produced phages are the P2 particles. Here, the consequences of these different strategies in the pirate and helper phage spreading among uninfected host is analyzed by using mathematical models. It is found that SaPI’s strategy interferes with the helper phage spreading significantly more than the P4’s strategy, because SaPI interferes with the helper phage’s main reproduction step, while P4 interferes only by forcing the helper lysogens to lyse. However, the interference is found to be weaker in the spatially structured environment than in the well-mixed environment. This is because, in the spatial setting, the system tends to self-organize so that the helper phages take over the front of propagation due to the need of helper phage for the pirate phage spreading.

**Competing interests:** The author declares no competing interest.

## Introduction

Spreading of pathogenic genes including antibiotic-resistance genes and toxin genes among bacteria is a real threat to the human society [1, 2]. One of the well-known examples of the mobile genetic elements that contribute to the spreading process is the *Staphylococcus aureus* pathogenicity islands (SaPIs). SaPIs spread among the host bacteria by using bacteriophages as their “helper” [3, 4, 5, 1, 2]. Interestingly, the mechanism of SaPIs spreading has many aspects in common with that of the well-studied pirate phage P4, that uses the helper phage P2 to spread among the host *Escherichia coli* [3, 6, 5].

The pirate phages and the helper phages we consider here all belong to the order *Caudovirales* or the tailed bacteriophages [7, 5], hence a phage particle consists of a head that contains the double-stranded DNA genomes and a tail. However, SaPI or pirate phage P4 does not have the structural genes that encode head and tail proteins, thus it cannot produce its progeny when infecting a host cell alone, and typically it becomes a prophage (Fig. 1a). Helper phages for them (e.g., *ϕ*11 for SaPIbov1, P2 for P4) are temperate phages. When infecting a sensitive host cell, it can enter either the lysogenic pathway with probability *α_H_* where the phage genome is integrated to the host chromosome as a prophage, provide immunity to the same phage, and replicate with the host, or the lytic pathway with probability 1 − *α_H_* to produce multiple copies of the phage particles and come out by lysing the host cell (Fig. 1b). Interestingly, pirate phages encode genes that allow them to interfere with the helper phage burst if it happens in the same host cell, including a factor to modify the size of helper head capsid so that it becomes too small to carry the helper genome. As a result, pirate phages take over the structural proteins produced from helper genes and produce pirate phages. This phenomenon was termed as “molecular piracy” [5].

**Figure 1:**
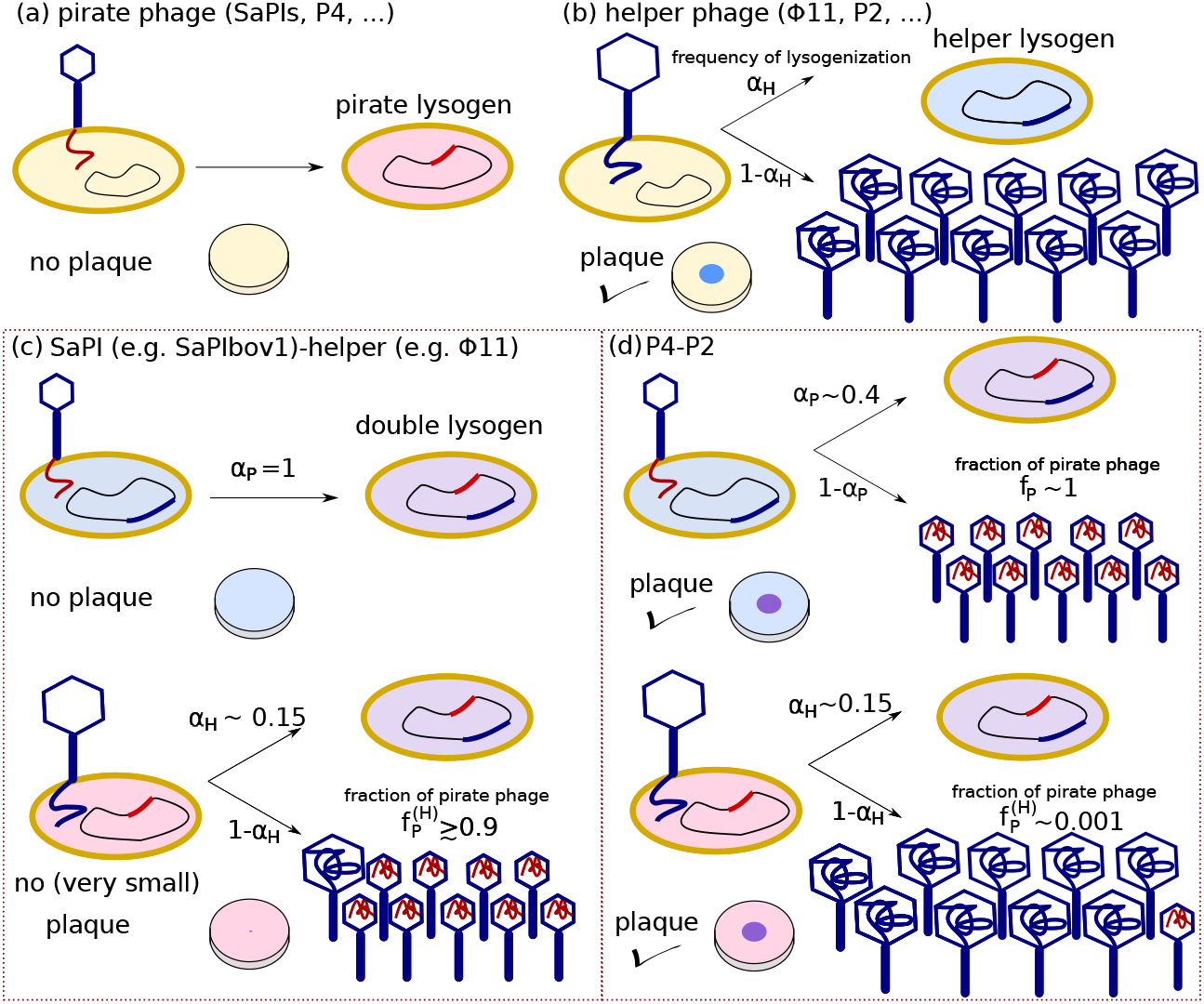
Strategies of SaPI-helper system and P4-P2 system. (a) Pirate phage can only become lysogen when infecting a host alone, hence it cannot propagate on a lawn of uninfected host. (b) Helper phage is temperate, with a frequency of lysogenization *α_H_*. Typically helper phage forms a turbid plaque on a lawn of uninfected host. (c) In the SaPI-helper system (e.g. SaPIbov1-*ϕ*11), infection of pirate phage SaPI particle to a helper lysogen results in formation of double lysogen (frequency of lysogenization of pirate phage upon infection of helper lysogen *α_P_* is one). Hence SaPI cannot propagate on a lawn of helper lysogen. When a pirate phage infects a SaPI lysogen and choose the lytic pathway with probability 1 −*α_H_*, the majority of the produced particles are pirate phages (fraction of pirate phage upon helper lysis decision 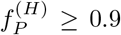), which inhibits the helper phage propagation on a pirate lysogen lawn. (d) In the P4-P2 system, infection of P4 phage to a P2 lysogen can results in the lysis with probability 1 − *α_P_* ~ 0.6, and upon burst almost all of the phages produced are P4 (fraction of pirate phage upon pirate lysis decision 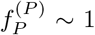), allowing pirate phage propagation on a helper lysogen lawn. On the other hand, P4 cannot take over much of P2 when P2 infection lyses a P4 lysogen (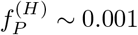), which allows helper phage propagation on a pirate lysogen lawn.

The fascinating mechanisms of the molecular piracy have been intensively studied, first for P4 which was found in 1963 [8]. SaPIs have attracted attention as a cause of the outbreak of toxic shock syndrome [9] and on-going studies are revealing the importance of them in *S. aureus* [2]. The pirate phages are ubiquitous; it has been found that the P4-type sequences are widespread among *E. coli* strains [10], while on average one SaPI was found per natural *S. aureus* strain [2]. These findings demonstrate the importance of the pirate phages as a source of horizontal gene transfer in bacteria.

When looking closer, however, SaPIs and P4 appear to have chosen very different strategies in their behaviour. In the case of SaPI, when it infects a helper lysogen, it simply integrates its genome to the host genome to form a double lysogen [11]. However, if a helper phage infects a SaPI lysogen and goes lytic (or the helper prophage in a double lysogen is induced), majority (fraction 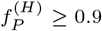) of the produced phage particles are SaPI particles [12]. The interference is so severe that helper phages are normally not able to make a visible plaque on a pirate lysogen lawn [4] (Fig. 1c). On the contrary, when infecting a helper P2 lysogen, a pirate P4 phage chooses lysogenic pathway with probability *α_P_* ~ 0.4, otherwise, it chooses lysis [13]. Upon bust, almost all the produced phages are P4 (fraction 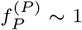) [14], i.e., P4 phage can spread on a lawn of P2 lysogen. When a P2 phage infects a P4 lysogen, the interference is weak, and upon lysis only small part (fraction 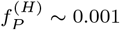) of the produced phages are P4 [15], enabling P2 phage to spread on a lawn of P4 lysogens (Fig. 1d). The parameters that illustrate these differences are summarized in Table 1.

**Table 1:** Key parameters for pirate phage strategies.

|  | Meaning | SaPI-helper | P4-P2 |
| --- | --- | --- | --- |
| $\alpha_P$ | Lysogen frequency upon pirate phage infection | 1 | 0.4 |
| $f_P^{(P)}$ | Fraction of pirate phage when the pirate phage lysing helper lysogen | N.A. | 1 |
| $f_P^{(H)}$ | Fraction of pirate phage when the helper phage lysing pirate lysogen | 0.9 | 0.001 |

Since both SaPIs and P4 are widespread, both choices seem to be viable. However, at least when looking at spreading on lysogens, the SaPI-helper system is inhibiting each other, while the P4-P2 system is supporting each other. A nontrivial question is how they spread when both pirate phage and helper phage encounter a population of host cells without any prophage. Can both the pirate and helper phages spread among the host, or does the interference prevent the spreading? What is the difference in the outcome between the two different strategies?

In order to answer these questions, we study the spreading of pirate phage and helper phage among uninfected host cells at a population level by using mathematical models. Previous studies on phage-bacteria systems have shown that the Lotka-Volterra type population dynamics model can reproduce many of the important phage-bacteria dynamics reasonably well in various length and time scales [16, 17, 18, 19, 20, 21, 22, 23, 24, 25, 26, 27, 28]. As a simple and well-defined setup, we here analyze the following two cases: Overnight growth in a culture of uninfected cells well-mixed with the pirate and helper phages, and the plaque formation in a lawn of uninfected cells initially spotted by a droplet of pirate-helper phage mixture. For the well-mixed case, we use the Lotka-Volterra type equations. For the plaque simulation, we let bacteria grow locally while letting phage and nutrient diffuse, describing the system using a reaction-diffusion type model [19, 26]. In both cases, we consider the time scale of an overnight, where the effect of the induction and the mutation are negligible. We compare the two phage strategies reflected in the values of 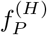, *α_P_*, and 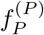, to reveal their impact on the phage spreading.

## Model

### Growth in a well-mixed culture

We use the Lotka-Volterra type ordinary differential equations to model the population dynamics of the system, which has been used to describe temperate phage and bacteria population dynamics [18, 23, 29, 26]. To extend the model to the pirate-helper phage system described in Fig. 1, we consider the follow-ing variables as functions of time *t*; uninfected bacteria concentration *B*(*t*), the pirate phage lysogen concentration *L_P_*(*t*), the helper phage lysogen concentration *L_H_*(*t*), the pirate phage concentration *P*(*t*), the helper phage concentration *H*(*t*), intermediate states concentration *I*(*t*) with subscripts that express each infection pathway, and the nutrient concentration *n*(*t*). Note that, in the case of P4, 99% of the case the genome is integrated into the chromosome, but 1% go into the plasmid state [13]. We here ignore these distinctions for simplicity and treat them as lysogens. We express these populations in the unit of number per volume.

When the lytic pathway is chosen upon infection, the free phages are produced after intermediate steps. Naturally, when a helper phage infects an uninfected bacterium and choose lysis, it will produce helper phages only as long as there is no superinfection by a pirate phage in the early part of the lytic pathway. The production of the pirate phage (or mixture of the helper and the pirate phages) can happen in different ways: (i) When a pirate phage infects a helper lysogen and force the lytic pathway with the probability 1 − *α_P_*. (ii) When a helper phage infects the pirate lysogen and choose the lytic pathway with probability 1 − *α_H_*. Further, we assume that some pirate phages can be produced (iii) when a pirate phage infects a bacterium in an early stage of the helper lytic pathway, to model the (almost) simultaneous co-infection. The model reflects these different possibilities for production of phages. The resulting model equations are as follows:

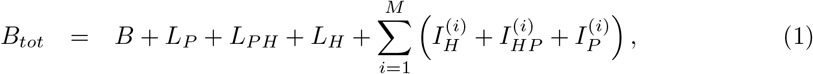

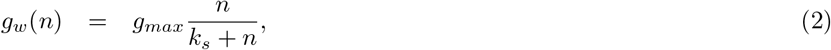

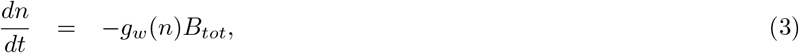

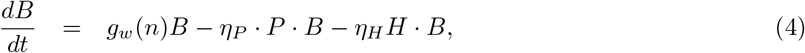

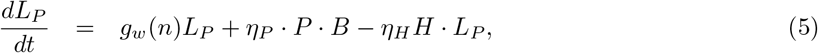

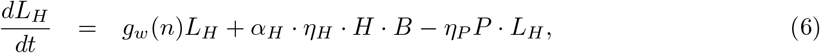

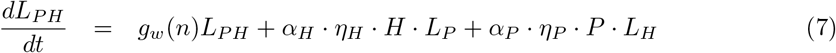

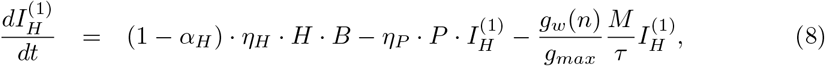

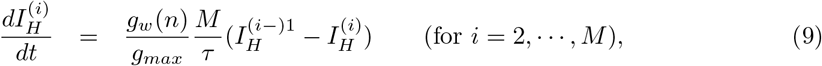

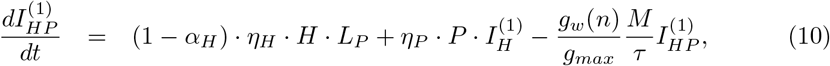

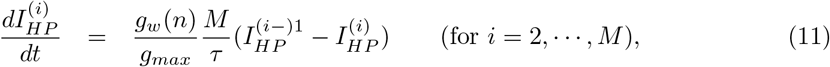

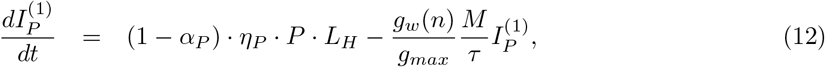

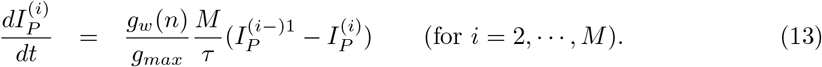

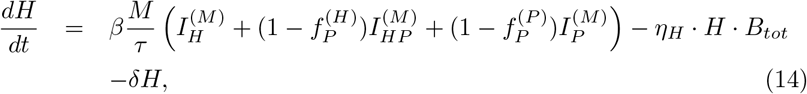

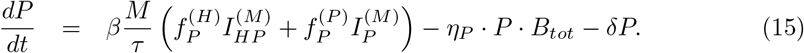

Here, we assume Monod’s growth law [30] with the maximum growth rate *g_max_* and the Michaels constant *k_s_*. Nutrient is measured in a unit where the yield for bacteria growth is one. We assume that the bacteria that are in intermediate steps of lysis consume the nutrient at the same rate as the growing bacteria. The latency time *τ* and the total number of phage particles produced per burst *β* are assumed to be the same for all infection pathways for simplicity. The intermediate infection steps are divided into *M* = 10 steps, which gives about 30% fluctuation in the latency time [26]. We assume that the latency time becomes longer proportional to the doubling time, ensuring that the phage production stops when the bacteria growth stops due to the lack of nutrient. Free phages decay at the rate of *δ*, but it is in general small [31] that it does not affect the present result.

### Initial condition and Numerical integration

We consider the population dynamics when the uninfected cells are mixed with the pirate-helper phage lysate in a rich medium. This is represented by initially having finite uninfected bacteria and nutrient as *B*(0) = *b*_0_ and *n*(0) = *n*_0_. The average phage inputs at time zero of helper phage *API_H_* and pirate phage *API_P_* determine the initial phage concentrations as *H*(0) = *b*_0_ · *API_H_* and and *P* (0) = *b*_0_ · *API_P_*.

The numerical integration was done by the 4-th order Runge-Kutta method, with time step 10^−6^/h.

### Plaque formation in a spatially structured environment

We model a plaque formation experiment setup called the spot assay [32], where a large number of bacteria in a soft agar is cast in a thin layer on a hard agar plate that contains the nutrient, a droplet of liquid that contains phages is placed and subsequently incubated overnight. With this method, it is easy to co-infect the bacteria with both helper and pirate phages in a well-controlled manner. Bacteria cannot swim visibly in the high viscosity of soft agar, hence each initially casted bacterium grow and divide locally to form a microcolony [33, 34]. As time goes, phages produced upon lysis of infected bacteria reach neighbour bacteria by diffusion to infect them and continue spreading. When the nutrients run out, both bacteria and phage growth ceases, leaving the plaque in the final configuration. When the phage is temperate, lysogens can grow in the infected area, resulting in a turbid plaque.

The clear plaque formation of virulent phage has been modeled by using reaction-diffusion type models [19, 35] and extended for turbid plaque formation by the temperate phage [26]. Here, we modify the turbid plaque formation model proposed in [26] to include the pirate phages. The latency time for the phage burst is taken into account by considering the intermediate states, and bacteria grows locally while the phage and the nutrient diffuse in space. Even though the local growth of bacteria into microcolonies may provide extra protection against phage attack [34, 27], for simplicity we here assume interaction in a locally mixed population. We use the same symbols as the well-mixed case to express the different populations, but now they are expressed in the unit of number per area by summing up the population over the thin soft-agar thickness Δ*a*, and they are a function of position **r** and time *t*. Furthermore, the diffusion terms are added for the nutrient and phages. The resulting model equations are parallel to the ones in eqs. (1)-(15), but the total derivative by time in the left-hand side of eqs. (3)-(15) should be replaced with partial derivative by time, and the diffusion terms *D_n_*∇^2^*n*, *D_H_* ∇^2^*H*, and *D_P_* ∇^2^*P* should be added to the right hand side of eqs. (3), (14), and (15), respectively. Furthermore, the pirate (helper) phage adsorption rate *η_P_* (*η_H_*) needs to be divided by Δ*a* reflecting that we are summing up the populations over this depth. Finally, to reflect the change of unit of the nutrient from per *ml* to per *μ*m^2^, the symbol for the Michaels constant is changed from *k_s_* in the well-mixed equation (2) to *K_s_* for the spatial model (used in the parameter table in Table 2).

**Table 2:** Default parameters used.

|  | Meaning | Default value | References, comments |
| --- | --- | --- | --- |
| $\eta_H$ | Helper phage adsorption rate | $3.3 \times 10^3 \mu\text{m}^3/\text{h}$ | P2 [31] |
| $\eta_P$ | Pirate phage adsorption rate | $13 \times 10^3 \mu\text{m}^3/\text{h}$ | P4 [31] |
| $\tau$ | latency time at fast growth | 0.75 h | P2 latency 30min[14]~ 48min [31]<br>P4 latency on P2 lysogen 60 min [14, 31] |
| $\beta$ | Total burst size | 100 | P2 on uninfected host: 60[45]~ 160[31]<br>P4 on P2 lysogen: 100[15]~300[31]<br>$\phi 11$ on uninfected host: ~100 [46] |
| $\alpha_H$ | Lysogenization frequency for a helper phage | 0.15 for both systems | P2: 0.15 ~ 0.2 [45]<br>$\phi 11$: ~ 0.14 [47] |
| $\alpha_P$ | Lysogenization frequency when a pirate phage infects a helper lysogen | 0.4 for P4-P2<br>1 for SaPI-helper | 0.3 ~ 0.5 [13]<br>1 [5] |
| $f_P^{(H)}$ | fraction of pirate phage from a helper phage lysing a pirate lysogen | 0.001 for P4-P2<br>0.9 for SaPI-helper | Yield is less than one per cell [15]<br>0.9-0.99 for SaPI-helper systems [12] |
| $f_P^{(P)}$ | fraction of pirate phage from a pirate phage lysing a helper lysogen | 1<br>(N.A. for SaPI) | $10^{-5} \sim 10^{-3}$ P2 and 100 P4 [14]<br>P2 level possibly due to the induction |
| $\delta$ | Phage decay rate | 0.001/h | Small in the time scale of interest. |
| $g_{max}$ | Fastest growth rate | 2/h | Typical *E. coli* growth in rich medium |
| $M$ | number of intermediate steps | 10 | 30% variation in latency time [26]. |
| $n_0$ | Initial nutrient level | $10^9$ /ml | well-mixed case |
| $k_s$ | Constant for Monod growth in the well-mixed simulation | $n_0/5$ | Depends on the nutrient. |
| $N_0$ | Initial nutrient level | $30/\mu\text{m}^2$ | spacially structured case [26] |
| $K_s$ | Constant for Monod growth in the plaque simulation | $N_0/5$ | Depends on the nutrient. |
| $D_H$ | Helper phage diffusion constant | $10^4 \mu\text{m}^2/\text{h}$ | 4-fold less than the value used for $\lambda$ [26]. $\lambda$ similar size to P2 [31] but $\lambda$ was selected for bigger plaque [48]. |
| $D_P$ | Pirate phage diffusion constant | $D_H \times \eta_P/\eta_H$ | The difference in the adsorption likely due to the difference in diffusion. |
| $D_n$ | Nutrient diffusion constant | $4 \times 10^5 \mu\text{m}^2/\text{h}$ | [26] |

### Initial condition and Numerical integration

We solved the model partial differential equations in polar coordinates with the origin at the center of the initially placed droplet. We assume the symmetry in a circular direction, and the spatial variation is only considered along the radial direction *r*. At a distant outer boundary *r* = *R_max_*, which is set to be 9 mm, we impose reflecting boundary condition for both phage and nutrient concentrations.

We analyze the plaque spreading in a lawn of uninfected bacteria. This is represented by initially distributing uninfected bacteria and nutrient uniformly as *B*(*r*, 0) = *B*_0_ and *n*(*r*, 0) = *N*_0_, with the rest of the variables set to zero, except for the phages placed in the middle where the phages are spotted. The average phage inputs in the initial spot of helper phage *API_H_* and pirate phage *API_P_* determine the initial phage concentrations. The initial spot of radius *R_s_* = 1mm is given by setting *H*(*r*, 0) = *B*_0_ · *API_H_* · Θ(*R_s_* − *r*) and *P* (*r*, 0) = *B*_0_ · *API_P_ ·* Θ(*R_s_* − *r*), where Θ(*x*) is the Heaviside step function. We set *B*_0_ =1/(18*μ*m)^2^ throughout this paper [26].

The integration was done by the finite difference method and time integration was done by the 4-th order Runge-Kutta method. The size of the spatial discretization, Δ*r*, was set to 18*μm*, hence at the initial condition, there is one bacterium per Δ*r*^2^ (*B*_0_ = 1/Δ*r*^2^). Time step of integration was set to Δ*t* = 10^−4^*h*.

### Parameters

Table 2 shows the list of the default parameters used, with the references for each value when available. Most of the values are chosen from the well-studied P4-P2 system, but when known the value for SaPI and a relatively well-studied helper phage for SaPIs, *ϕ*11, are also stated with references. The difference in the strategies are reflected in the values of 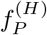, *α_P_*, and 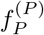, and the rest of the parameters are kept same between the two systems.

## Results

### Growth in a well-mixed culture

In order to compare the SaPI-helper strategy and the P4-P2 strategy, we set most of the parameters same for both systems as described in Table 2. The difference in strategy is reflected by choice of the following parameters as summarized in Table 1: the pirate frequency of lysogeny upon infection of helper lysogen *α_P_*, the fraction of the pirate phage production upon a pirate phage lysing a helper lysogen 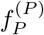, and the fraction of the pirate phage production upon a helper phage lysing a pirate lysogen 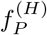. The detail of the model equations are given in the Model section.

We first simulate the case where uninfected cells of concentration *b*_0_ = 10^5^/ml are mixed with the pirate phage and the helper phage lysate. We vary the initial average phage input for the pirate phage *API_P_* and that for the helper phage *API_H_*, by mixing *API_P_* · *b*_0_ pirate phages and *API_H_* · *b*_0_ helper phages to the culture at time zero. As the bacteria cells grow, the nutrient in the culture is consumed. The initial nutrient concentration was set so that the population would reach 10^9^/ml if there were no killing by phage. When the nutrient concentrations become zero, both the bacteria and the phage stop growing, though free phage adsorption to the bacteria still happens.

Figures 2 show the time course of the population concentrations for the SaPI-helper strategy (Figs. 2a) and the P4-P2 strategy (Figs. 2b), with *API_P_* = *API_H_* = 100.

**Figure 2:**
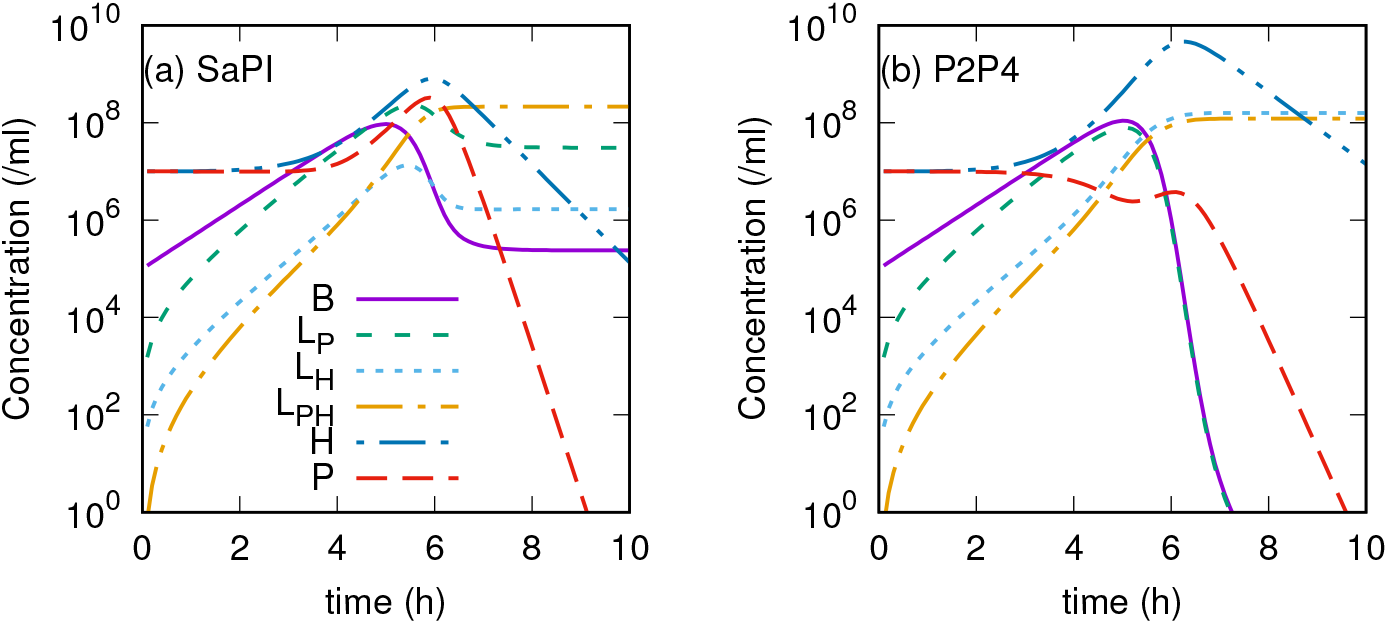
Simulated well-mixed population dynamics with (a) SaPI-helper (b) P4-P2 system. The initial condition is set so that *API_P_* = *API_H_* = 100. The concentrations of sensitive bacteria *B*, the pirate lysogen *Lp*, the helper lysogen *L_H_*, the double lysogen *L_PH_*, the helper phage *H*, and the pirate phage *P* are shown by solid line, coarse dashed line, fine dashed line, dash-dotted line, dash-double dotted line, and long dashed line, respectively.

In the case of the SaPI-helper system, we can see that the pirate phage concentration *P* rises following closely with the increase of the helper phage concentration *H*. The delay is because the pirate phage can replicate only if the pirate lysogens are infected by the helper phage, while helper phage can just replicate upon single infection. At first, the initially added pirate phages typically form lysogens and they grow, as reflected in the increase of *L_P_*. The replicated helper phage starts to infect these pirate lysogens, and they produce mainly the pirate phages. This pirate phage replication severely interfere with the helper phage replication, hence overall phage replication rate is low (compare with the P4-P2 strategy Figs. 2b). As a result, a significant number of bacteria are left uninfected when the nutrient is depleted.

This is in contrast to the P4-P2 system, where the pirate phage can take over the helper phage only when it infects a helper lysogen. For the increase of the pirate phage, the increase of the helper lysogen *L_H_* needs to happen first, which allows the helper phages to grow without feeling the interference by the pirate phage for quite some time. The pirate phage *P* starts to increase only about 6h after the start of the simulation, but the system soon runs out of the nutrient and thus pirate phages do not increase much. At the same time, the lack of interference to the helper phage replication allows the helper phages to infect all the uninfected cells *B* and the pirate phage *L_P_*, converting them to either helper lysogens *L_H_* or the double lysogens *L_PH_*.

### Initial phage input dependence of the prophage spreading in well-mixed environment

The degree of interference between the helper phage and the pirate phage should depend on the initial phage inputs. This is studied in the Figs. 3 for each strategy. Since the free phages are eventually adsorbed in the bacteria, we consider the number of the lysogens with helper prophage *L_H_* + *L_PH_* in the final state as a measure of the helper phage spreading, while the lysogens with pirate prophage *L_P_* + *L_PH_* as a measure of the pirate phage spreading.

**Figure 3:**
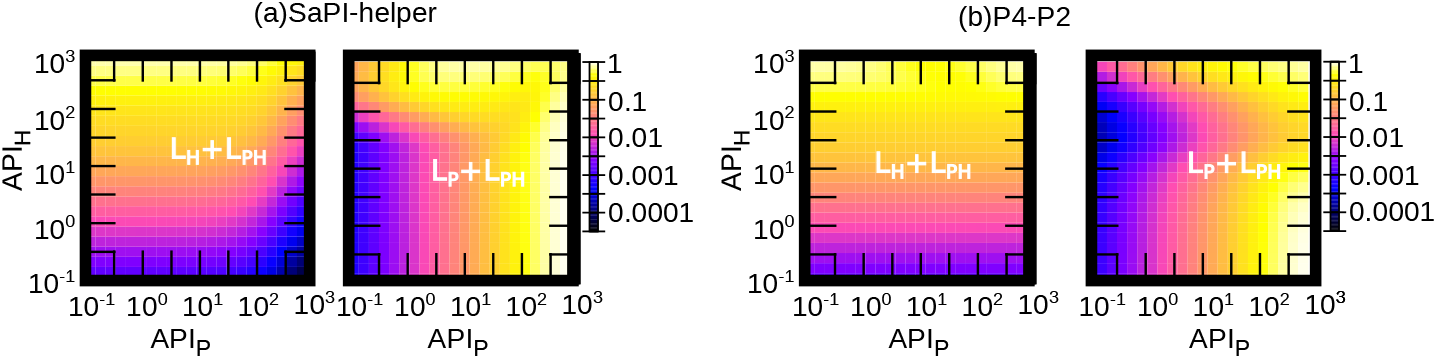
The initial condition dependence of the helper and pirate phage gene spreading in a well-mixed population. In the initial condition, the uninfected bacteria of 105/ml is mixed with the phage particles of various average phage input *API_H_* and *API_P_* and the population dynamics was simulated until the system reaches the final state by using (a) the SaPI-helper system’s parameter set (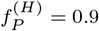, *α*_*P*_ = 1) or (b) the P4-P2 system’s parameter set (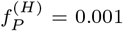, *α_P_* = 0.4, 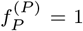). The number of lysogen cells with helper prophage *P P L_H_* + *L_PH_* (pirate prophage *L_P_* + *L_PH_*) in the final state normalized to the initial nutrient concentration 109/ml are visualized by color code in figure left (right) panel, respectively, where the higher value is shown with lighter color in log scale.

Clearly, in the SaPI-helper system, a larger amount of the pirate phage in the initial culture (large *API_P_*) interferes more severe with the helper phage spreading (Fig. 3a left), especially when the helper phage input *API_H_* is relatively small. However, when *API_P_* is high, the pirate phage spreading is not compromised much from the lack of helper phage growth (Fig. 3a right), since the initial amount of pirate phage is already high enough and the formed pirate phage lysogens can grow over time. At the same time, the pirate phage clearly benefit from taking over the helper phage for replication when *API_P_* is low but *API_H_* is high, reflected in the increased amount of *L_P_* + *L_PH_* when increasing *API_H_* with keeping *API_P_* ~ 0.1.

On the contrary, in the P4-P2 system, the final concentration of bacteria with helper prophage *L_H_* + *L_PH_* is almost independent of *API_P_* (Fig. 3a right), indicating that the helper phage spreading is not interfered by the pirate phage much. Interestingly, the pirate phage spreading at a fixed *API_P_* showed a non-monotonic dependence on *API_H_*. This is because the lysis of the pirate lysogen by helper infection does not contribute to the pirate phage production, and the pirate phage needs to lyse helper lysogen for pirate phage production; increasing *API_H_* for a fixed *API_P_* contribute both processes while the latter become dominant only when *API_H_* is high enough so that initially added pirate phages infect the helper lysogens and not the uninfected host. When *API_H_* value is moderate, it can lyse the pirate lysogens formed due to initially added pirate phages, which reduces the amount of pirate lysogens.

When comparing the two strategies, SaPIs are better at spreading than P4 in a well-mixed system, as depicted by the larger population of *L_P_* + *L_PH_* in all the studied parameter regions. At the same time, the helper phages are interfered strongly by SaPI but not by P4.

### Plaque formation in a lawn of uninfected bacteria

The analysis of the well-mixed system showed that SaPI significantly interferes with the helper spreading, while P4 does not. The strong interference by SaPI could make the spreading of SaPI in the spatially structured environment harder since if the helper phage is locally depleted by interference, the phages cannot spatially propagate even if there are uninfected cells outside of the plaque.

In order to see if the local depletion can be a problem, we next analyzed the plaque spreading in a lawn of uninfected bacteria. The simulation mimics the plaque formation by the spot assay [32], where soft agar with bacteria is cast over hard agar with nutrient, a droplet with mixture of phages is placed in the center, and incubated overnight. In the setup, bacteria cannot swim visibly and the local growth results in formation of a microcolony at the position where a bacterium was initially dispersed [33, 34]. Phages grow and diffuse to infect new bacteria, resulting in a circular zone of killing around the initially spotted area. When the phage is temperate, lysogens will survive and grow, making the plaque turbid. The simulation was started by placing a droplet of mixture of pirate and helper phage with radius 1mm. The concentration of phages in the droplet is defined by the average phage input per initially placed bacterium for the helper phage, *API_H_*, and that for the pirate phage, *API_P_*. The detail of the simulation is given in Model section.

The time courses of the plaque formation with *API_H_* = *API_P_* = 1 (i.e., on average there is one of each phage per initially placed bacterium under the initial spot) are shown in Fig. 4a-f for the SaPI-helper system and in Fig. 4g-l for the P4-P2 system, respectively. The populations at each location are shown as a function of distance from the infection origin, *r*. The unit of the population density is normalized to initial bacteria density *B*_0_.

**Figure 4:**
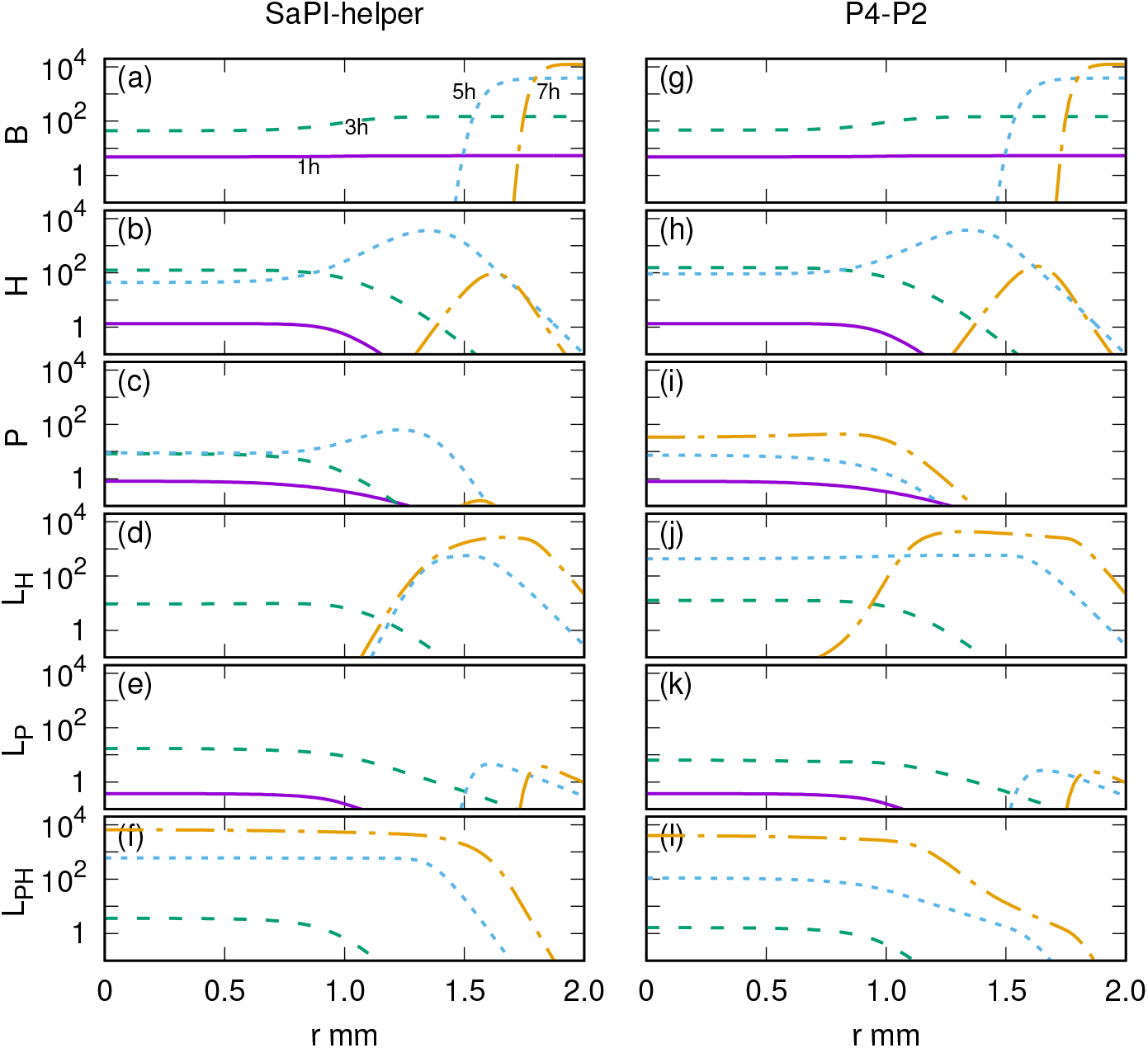
Simulated plaque formation by (a-f) SaPI-helper and (g-l) P4-P2 system. (a-f) and (g-l) represents the time evolution of the populations. The unit is the number of the cells/phage particles per 18*μm*^2^, corresponding to the number per microcolony. The profiles at 1h, 3h, 5h, and 7h are shown by solid line, coarse dashed line, fine dashed line, and dash-dotted line, respectively.

The profile for the uninfected bacteria and the helper phage at the front of the plaque spreading looks very similar between the two systems (Fig 4ab vs. gh). This is because, the helper phage properties are set to be identical when there is no pirate phage, and at the front of the plaque propagation, the helper phages does not feel the interference by the pirate phage due to their low density.

The difference can be seen in how the pirate phage can propagate. We see that pirate phage can come up much later and close to the center in the P4-P2 system (Fig. 4i), where pirate phages can be produced only when the helper lysogen *L_H_* is lysed by the pirate phage infection. In the SaPI-helper case (Fig. 4c), the wave of pirate phage follows closer to the front propagation of helper phage by taking over the part of the helper phage lysis, making it possible for the pirate phage to spread a lot faster and further away, producing larger amount of the double lysogen in the final state (Fig. 4f vs. l).

These results depict that the SaPI-helper strategy is indeed beneficial for the spreading of the pirate phage prophage spreading in space because the strong interference does not inhibit helper spreading as long as the helper phage takes over the front of the plaque propagation. The P4-P2 strategy, on the other hand, does allow the helper phages to spread, but the pirate phage gene spread is delayed more because pirate phage can grow only later, after the helper lysogens are established.

### Initial phage input dependence of the prophage spreading in spatially structured environment

The spreading of the phage gene should depend on the initial concentration of each phage, *API_H_* and *API_P_*, in the spot. When the initial pirate phage input *API_P_* is too high compared to *API_H_*, the strong interference of SaPI-helper strategy may inhibit spreading of both of the phages, while when *API_H_* ≫ *API_P_*, the P4-P2 strategy may better to spread since it can lyse the helper lysogens.

For systematic understanding, we investigated how the outcome changes when varying the initial concentration of the phages in the spot, *API_H_* and *API_P_*. The results are summarized in Fig. 5 for (a) the SaPI-helper strategy and (b) the P4-P2 strategy by the final population with the helper phage gene *L_H_* + *L_PH_* and the pirate phage gene *L_P_* + *L_PH_* integrated over space. The integration is done outside of the initial spot radius, in order to focus on the effect of the spreading.

**Figure 5:**
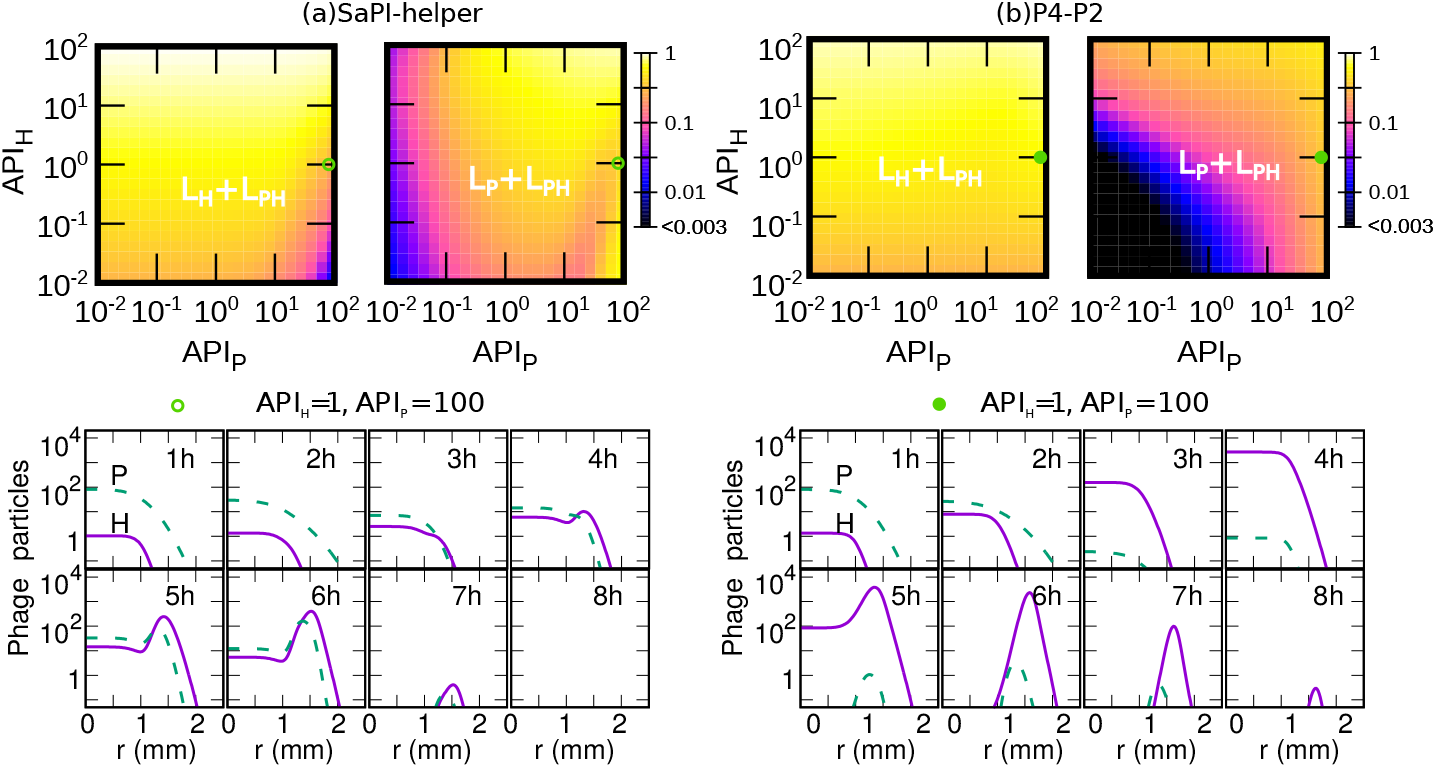
The initial condition dependence of the helper and pirate phage gene spreading. The average phage input in the initial spot, *API*_*H*_ and *API*_*P*_ is varied, and the plaque formation is simulated by using (a) the SaPI-helper system's parameter set (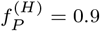, *α_P_* = 1) or (b) the P4-P2 system’s parameter set (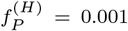, *α_P_* = 0.4, 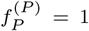). The number of lysogen cells with helper prophage *L_H_* + *L_PH_* (pirate prophage *L_P_* + *L_PH_*) integrated over the space outside of the initial spot (*r* > 1mm) and normalized to the maximum observed value 1 × 10^8^ are visualized by color code in figure top left panels (top right panels), respectively, where the higher value is shown with lighter color. The bottom panels show the time evolution of the helper phage (solid line) and the pirate phage (dashed line) particles propagation for the *API_H_* = 1 and *API_P_* = 100 case.

For the helper phage spreading, we observed that the interference by SaPI is relatively weak (Fig. 5a top left). This is because the system tends to self-organize so that the helper take over the spreading front. Once the helper takes over the front, the pirate phage cannot lyse the helper lysogen, hence there will be a large amount of helper prophages left. An example of such self-organization is depicted in the case of *API_H_* = 1 and *API_p_* = 100 by showing the time course of the pirate phage and helper phage profile over time (Fig. 5a bottom). At the start, the pirate phage is in the front of propagation, but it goes down over time since the pirate phages simply diffuse and get adsorbed. The helper phage can reproduce in small amount by infecting uninfected host, and at about 3h the helper phage catches up the propagation front. Then the helper phages start to replicate fast by lysing the uninfected cells at the propagation front, producing a peak of helper phage at 4h. This peak propagates with pirate phage peak closely following behind, because some of the helper phages lyses the pirate phage lysogens that were produced by the initially added pirate phages. Of course, this self-organization cannot happen when *API_H_* is too small and *API_P_* is too large, depicted by the significantly low level of helper phage spreading at *API_H_* ~ 0.01 and *API_P_* ~ 100. Overall, the pirate phage spread fairly well (Fig. 5a top right), because if the helper phage takes over the propagating front, pirate phage can follow closely, while when the initial pirate phage level is so high to completely inhibit helper phage propagation, the abundant initial pirate phages diffuses and produce the pirate lysogens.

In the P4-P2 system, again the helper phage spreading is not interfered by the pirate phage P4 (Fig. 5b top left), because even when the *API_P_* ≫ *API_H_* at first, the helper phages simply lyse the pirate lysogens to produce helper phage and propagate, as depicted in the very rapid take over of the front by helper phages in the *API_H_* = 1 and *API_P_* = 100 case shown in Fig. 5b bottom. Overall, the pirate phage spreading (Fig. 5b top right) increase with *API_P_* and *API_H_*, but the absolute level of final spreading of the pirate phage gene *L_P_* + *L_PH_* is significantly less than the SaPI case.

## Discussion

The pirate-helper phage system demonstrates the fascinating richness of the interaction between phages in nature. At the same time, given the increased number of possible infection scenarios, it becomes significantly complex to analyze the population dynamics. Still, the SaPI-helper system and the P4-P2 system pose interesting examples, in that their spreading among the lysogen populations are completely opposite (Fig. 1). The present analysis of spreading among the initially uninfected population showed that, in both well-mixed and spatially structured cases, SaPI is spreading better than P4. At the same time, SaPI strongly interferes with the helper phage spreading, while P4 does not interfere much with the P2 spreading. This is natural because SaPI takes over most of the phage production when the helper phage trigger the lysis (i.e., 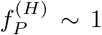), which is the major part of the helper phage production. On the contrary, even if the pirate phages can lyse the helper lysogens and produce mostly the pirate phage (*α_P_* < 1, 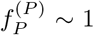) as P4 does, for helper phage that is some reduction of the existing helper lysogens but not the interference to the helper phage reproduction, hence effect is weak.

Another point that the present analysis has clarified is the weaker trade-off between the helper phage spreading and the pirate phage spreading in the spatially structured lawn of uninfected cells. Even if the interference by pirate phage upon helper burst is strong as 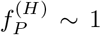, as long as the helper phages come to the front of spreading this does not inhibit the helper phage spreading, and pirate phage can spread as well by following the front. Because the pirate phage cannot spread by themselves, it is relatively easy for the helper phage to take over the front. The pirate-behind-helper configuration makes the most of the pirate phage lysogens be double lysogens with the helper. However, this contradicts somewhat to the observation that very few SaPIs are found as a double lysogen with its own helper phage [2]. The scarcity of SaPI-helper double lysogen could be due to strong selection pressure against being a helper phage because SaPI is inhibiting the helper phage so much upon induction of the double lysogen [2]. This possibility can be analyzed by extending the present population dynamics model to include inductions and mutations and simulate a longer time scale. Also, the present results relies on the assumption that bacteria do not migrate, hence the propagation of the phages relies only on the diffusion on top of the replication. The virulent phage propagation on the migrating population of bacteria has been found to show very rich patterns [36], and it will be interesting to study similar setup with pirate and helper phages.

It is an interesting and unsolved question why SaPI-helper system and P4-P2 system have such different strategies. It appears that SaPIs are spreading better than P4 at the expense of stronger interference with helper phages. From the pirate phage’s point of view, SaPI strategy appears better, but it is also possible that in the long run, the P4 strategy is more sustainable because the helper phage, which is a necessary host for the pirate phage, also grows well.

The pirate-helper phage interaction is formed under co-evolution of them, but in addition, other players are likely to contribute to the co-evolution. Naturally, the pirate-helper interaction parameters vary depending on the combination. The frequency of lysogenization *α_H_* for phage 80*α*, a helper phage for SaPIs, was found to be quite low (10^−5^ ~ 10^−3^) [37], which will change the outcome even if other parameters were identical. The frequency of lysogenization for helper phage *α_H_* may be selected mainly by the frequency of the host population collapse [38] or by competition with other temperate phages [29], independent of the interaction with the pirate phage, even though the value of *α_H_* do affect the outcome of the pirate-helper system strongly. More interestingly, the P4 phage cannot lyse the phage 186 lysogen, i.e., *α_P_* = 1, while interference upon the phage 186 bursting P4 lysogen is moderately strong (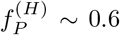 to 0.9)[39], which appear to be closer to the SaPI-helper strategy. It is difficult to tell how often a given pirate phage uses the found helper phage in natural environments, even though that could affect how much the system had co-evolved. It should also be noted that using multiple helper phages is beneficial by it-self since it greatly increases the host range of the pirate phages. However, at the same time, the selection for the ability for a pirate phage to interact with multiple helper phages may put some limit in optimizing the interaction with a specific helper phage due to the molecular differences of helper phages. In addition, the pirate phage may deliver the gene beneficial to the host bacteria as SaPIs carry pathogenic genes, and this can significantly contribute to spreading of the pirate phage gene, too.

Finally, the similarity and the differences of the pirate phages from the defective interfering particles (DIPs) in viral systems should be commented. DIPs are the defective viral particles that cannot replicate by itself upon infection of the host, but when co-infected the host with intact viruses, it will interfere with the production of intact viruses and produce DIPs instead [40, 41]. The known naturally occurring examples of DIPs such as those found in influenza virus are produced as the original virus replicates [40], but there are several efforts to engineer DIPs to interfere the spreading of pathogenic virus, including the HIV virus [42]. The noticeable difference between DIPs and pirate phage discussed in this paper is that the “helper” viruses for DIPs typically show chronic infection, i.e., infected host cells stay alive for a relatively long time and keep producing the viruses, and hence the helper virus does not have a dormant mode that corresponds to lysogeny in the helper temperate phage. As a consequence, there is no corresponding difference of the strategies of pirate phage replication that depends on the which phage determines to “go lytic”. Nevertheless, some of the population dynamics discussed here has some similarity to those of DIPs. For example, a recent theoretical paper that analyzed the spreading of the DIPs in a spatially structured environment [43] pointed out that the spreading speed of the virus is not affected by the DIPs if the virus takes the front of propagation, even if the wave of DIPs is propagating just behind. This is consistent with the present finding that once the helper phage takes over the front that is stably maintained (Fig. 5), and it is expected that also in DIP-virus system, there is a tendency to self-organize so that the intact virus take over the front. It is worth mentioning that experiment on the spreading of DIPs and viruses mixture in space reported highly heterogeneous pattern in some conditions [44], demonstrating the importance of stochasticity. It is an interesting future work to extend the model to include stochastic population fluctuations in pirate-helper phage system, as well as analyzing it experimentally.

Overall, this paper sheds light on one of the many facets of the art of war among bacteria and phages. Considering the ubiquity of the pirate phages, their impact on the ecology and evolution of the microbial system should be significant. Clearly, further study is required to understand the rich population dynamics of pirate-helper systems.

## Acknowledgments

NM thanks Kim Sneppen for fruitful discussions. This work was funded by the Danish National Research Foundation (BASP: DNRF120).

